# The Microbiota Dictates Vendor-Derived Differences in a Murine *Clostridioides difficile* Infection Model

**DOI:** 10.64898/2026.08.31.748273

**Authors:** Liam Keogh, Lharbi Dridi, Bavanitha Thurairajah, Michael Shamash, Corinne Maurice, Irah L. King, Bastien Castagner

**Affiliations:** Department of Pharmacology & Therapeutics, McGill University, Montreal, QC, Canada; Meakins-Christie Laboratories, Research Institute of McGill University Health Centre, Montreal, QC, Canada; Department of Microbiology & Immunology, McGill University, Montreal, QC, Canada; McGill Centre for Microbiome Research, Montreal, QC, Canada; McGill Regenerative Medicine Network, Montreal, QC, Canada; McGill Interdisciplinary Initiative in Infection and Immunity, Montreal, QC, Canada

## Abstract

*Clostridioides difficile* infection (CDI) is the leading cause of healthcare-associated infectious diarrhea and remains a major burden to healthcare systems worldwide. The development of novel therapeutics for CDI requires robust and reproducible preclinical models. However, the microbiota has emerged as a major source of variability in animal studies. Here, we found that genetically similar mice obtained from two commercial vendors, Jackson Laboratory (JAX) and Charles River Laboratories (CRL), exhibited marked differences in susceptibility to CDI, with JAX mice developing fulminant disease and CRL mice remaining resistant. Using full-length 16S rRNA gene sequencing, we show that JAX and CRL mice harboured distinct gut microbiota, and that cohousing susceptible JAX mice with resistant CRL mice was sufficient to shift the JAX microbiota toward the CRL community structure and confer resistance to CDI. Differential abundance analysis identified taxa distinguishing resistant and susceptible mice, providing candidates for future mechanistic investigation. These findings demonstrate that vendor-derived variation in the gut microbiota drives differential susceptibility to CDI in mice, and that this phenotype is transferable via cohousing, highlighting the importance of accounting for the microbiota when designing and interpreting animal models of infectious disease.

**IMPORTANCE:** Mice are widely used to study *Clostridioides difficile* infection, but animals purchased from different commercial vendors can respond differently to infection. This study shows that mice from two commonly used suppliers differ substantially in their susceptibility to *C. difficile*, and that this difference is caused by variation in their gut microbiota, as opposed to genetic differences. By housing susceptible and resistant mice together, we found that the protective bacterial community could be transferred to susceptible mice, making them resistant to severe infection. This work underscores that the choice of animal vendor is a critical experimental variable that can substantially influence infection outcomes, highlighting the importance of accounting for vendor source when designing, interpreting, and comparing mouse studies of *C. difficile* infection.

## INTRODUCTION

*Clostridioides difficile* is a Gram-positive, spore-forming, anaerobic bacterium that colonizes the gastrointestinal tract of humans and animals.^1, 2^ It is recognized as the leading cause of healthcare-associated infectious diarrhea and remains a major burden on healthcare systems worldwide.^3^ Although associated with hospitalization and antibiotic use, increasing rates of community-acquired *C. difficile* infection (CDI) have been reported, highlighting the public health significance of this pathogen.^4^

A central feature of CDI susceptibility is the disruption of the gut microbiota.^5^ In healthy individuals, the intestinal microbiota provides colonization resistance against *C. difficile* through multiple mechanisms, including nutrient competition, production of antimicrobial metabolites, secondary bile acid metabolism, and modulation of host immune response.^6^ Antibiotic exposure is the most common cause of CDI as it disrupts these protective microbial communities.^2^ The resulting loss of microbial diversity and functional capacity creates a permissive niche for *C. difficile* spore germination, proliferation, and toxin production.^2, 7^ Specific microbial taxa and metabolic functions within the gut microbiota have been shown to strongly influence susceptibility to both *C. difficile* colonization and toxin-mediated damage.^8-12^ Despite the central role of antibiotic-induced dysbiosis in CDI pathogenesis, first-line treatment remains vancomycin and fidaxomicin.^13^ These therapies often fail to restore a protective microbiota and may further prolong dysbiosis.^14^ As a result, 15-25% of patients experience recurrent infections after initial treatment.^13^ This underscores the need for therapeutic approaches that move beyond antibiotics and instead target key aspects of the pathogen life cycle. Emerging strategies include neutralizing toxin activity, inhibiting spore germination, and microbiome-directed interventions aimed at restoring colonization resistance.^15, 16^ However, the development and evaluation of these novel interventions require robust and reproducible animal models to study CDI.

Several animal models have been developed to study CDI, including mouse and hamster models.^17-20^ Among these, mouse models are most widely used, encompassing a range of disease phenotypes, from asymptomatic colonization to severe infection.^17^ A commonly used approach is described by Chen et al., in which conventional C57BL/6 mice are administered an antibiotic cocktail prior to *C. difficile* challenge.^21^ This regimen strongly disrupts the gut microbiota and typically results in severe disease, with mortality occurring within the first week following challenge. Owing to its robust disease phenotype, this model has become widely used for evaluating candidate therapeutics and investigating mechanisms of pathogenesis.^22-25^

An important but often overlooked consideration in these models is the source of experimental animals. The C57BL/6 mouse is the most widely used strain is CDI research.^17^ However, animals obtained from different commercial vendors can differ substantially in both genetic background and gut microbiota composition.^26, 27^ Genetic drift, for example, has produced known genomic differences between C57BL/6J from the Jackson Laboratory (JAX) and C57BL6/N mice from Charles River Laboratories (CRL).^28^ Vendor-specific microbial communities can also persist even after animals are transferred to a shared housing environment, and such differences have been shown to alter immune responses, metabolism, and susceptibility to infectious and inflammatory disease across multiple murine models.^27, 29-32^

Despite the widespread use of C57BL/6 mice in CDI research, little attention has been given to the potential influence of vendor-specific microbiota differences on experimental outcomes. It remains unclear differences in baseline microbiota composition affect susceptibility to fulminant CDI, or whether these differences are driven primarily by the microbiota rather than host genetics. Such vendor-associated microbiota variation could contribute to variability across studies and confound the interpretation of therapeutic efficacy.

The objective of this study was to determine whether C57BL/6 mice obtained from two commonly used vendors, The Jackson Laboratory and Charles River Laboratories, differ in their susceptibility to *C. difficile* challenge. After identifying differences in disease severity between vendors, we investigated whether this was microbiota-dependent by cohousing mice from both vendors to facilitate microbial exchange. We then performed 16S rRNA gene sequencing to characterize baseline differences in microbiota composition between vendors and assess how cohousing and antibiotic treatment altered the microbiota over time. By defining the contribution of vendor-associated microbiota to CDI susceptibility, this study aims to improve the reproducibility and interpretation of murine CDI models and identify microbial taxa that may influence disease outcome.

## RESULTS

### JAX and CRL mice have different susceptibility to *C. difficile* challenge

To determine how vendor origin influences the susceptibility to *C. difficile* challenge, we obtained two substrains of C57BL/6 mice from vendors commonly used for CDI studies: C57BL/6J (JAX) from The Jackson Laboratory, and C57BL/6N (CRL) from Charles River Laboratories. The mice were treated with the antibiotic model described by Chen et al., followed by challenge with 1x10^5^, 10x10^6^, or 1x10^7^ CFU *C. difficile* VPI 10463 spores (Figure 1A).

**Figure 1:**
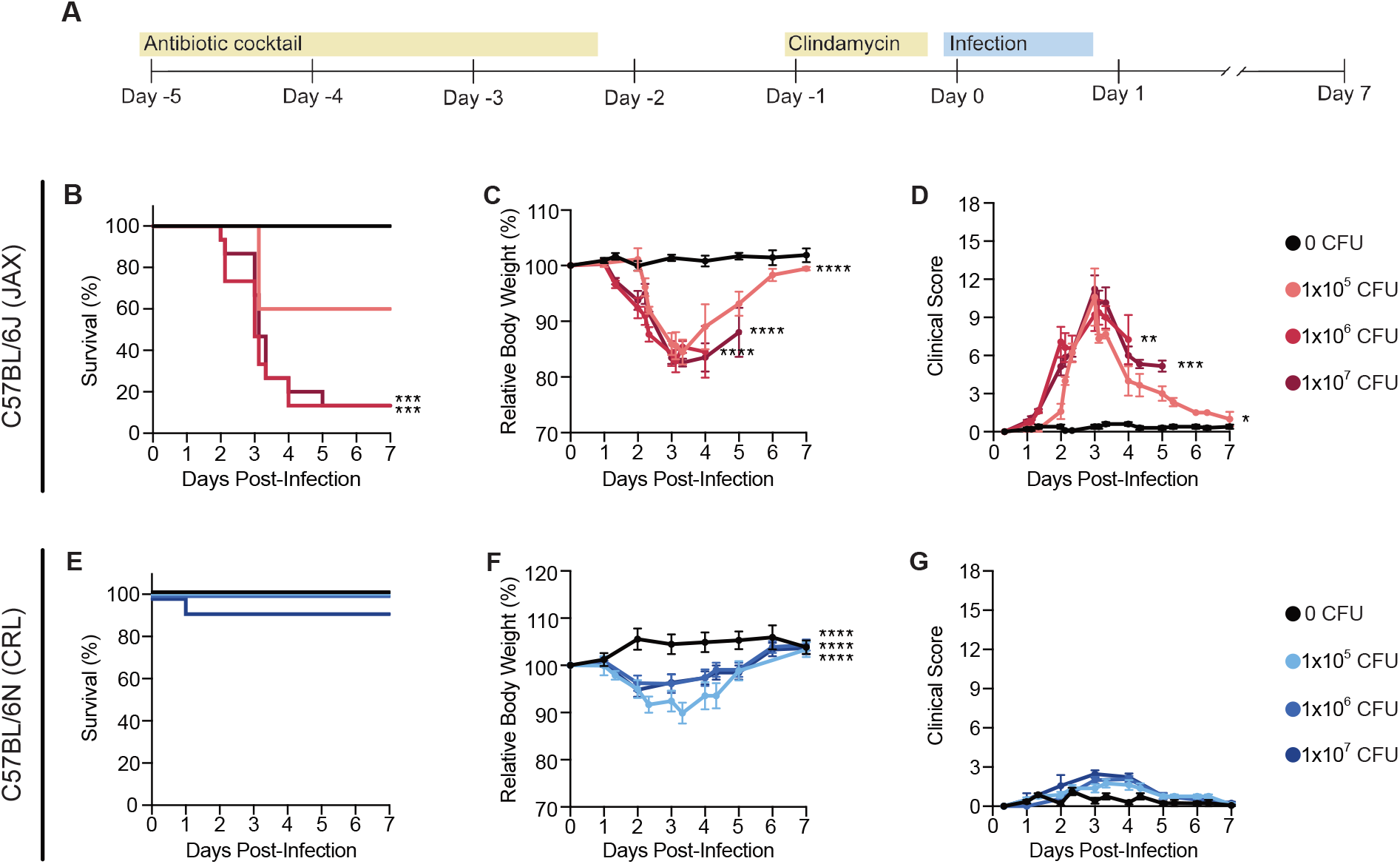
Vendor-dependent differences in susceptibility to *C. difficile* challenge: (A) Experimental design: mice received an antibiotic cocktail and clindamycin to induce dysbiosis, followed by *C. difficile* challenge with 0 (uninfected control), 1x10^5^, 1x10^6^, or 1x10^7^ CFU spores, and were monitored for 1-week post-infection. (B-D): JAX mice: (B) Kaplan-Meier survival, (C) relative body weight (% of baseline at Day 0), and (D) disease severity. (E-G): CRL mice: (E) Kaplan-Meier survival, (F) relative body weight (% of baseline at infection), and (G) disease severity. Panels C, D, F, and G show mean ± SEM. \**P* ≤ 0.05; \*\**P* ≤ *0.01;* \*\*\**P* ≤ *0.001;* \*\*\*\**P* ≤ *0.0001* (comparisons to uninfected control). n = 6-14 mice per group (See Fig. S1).

*C. difficile* burden was comparable between JAX and CRL mice during the first 5 days post-infection (*P* = 0.057; Figure S1C), indicating that both groups were equally permissive to *C. difficile* colonization and outgrowth. Disease outcomes, however, diverged substantially despite comparable colonization. JAX mice developed fulminant disease, with mortality occurring between days 2 and 5 post-infection. At both the inocula of 1x10^6^ and 1x10^7^ CFU, mortality reached 87% (13/15), with a median survival time of 3 days post-infection (Figure 1B). Consistent with this mortality, JAX mice exhibited significant weight loss beginning at day 2 post-infection (Figure 1C), coinciding with the peak of disease severity, characterized by diarrhea, reduced activity, ruffled coat, and ocular/nasal discharge (Figure 1D; S1A).

In contrast, CRL mice were largely resistant to severe CDI. Across all inocula tested, mortality did not differ significantly from uninfected controls (Figure 1E). Although CRL mice exhibited modest weight loss relative to uninfected controls (Figure 1F), overall disease severity was reduced relative to JAX mice, and clinical scores in infected CRL mice did not differ from uninfected controls (Figure 1G). Clinical signs in CRL mice were generally limited to mild diarrhea and transient weight loss, and most animals recovered fully during the observation period (Figure S1B).

Together, these findings indicate that the divergence in CDI severity between JAX and CRL mice occurs independent of initial colonization efficiency, suggesting host- or microbiota-dependent factors associated with vendor origin as drivers of disease severity.

### Cohousing JAX and CRL mice alters the susceptibility to *C. difficile* challenge

To determine whether the different susceptibility between JAX and CRL mice was microbiota-dependent, we cohoused mice from both vendors prior to infection to facilitate microbial exchange. JAX and CRL mice were housed separately by vendor or cohoused in a 1:1 ratio (MIX) for 14 days prior to continuing antibiotic treatment and *C. difficile* challenge (Figure 2A).

**Figure 2:**
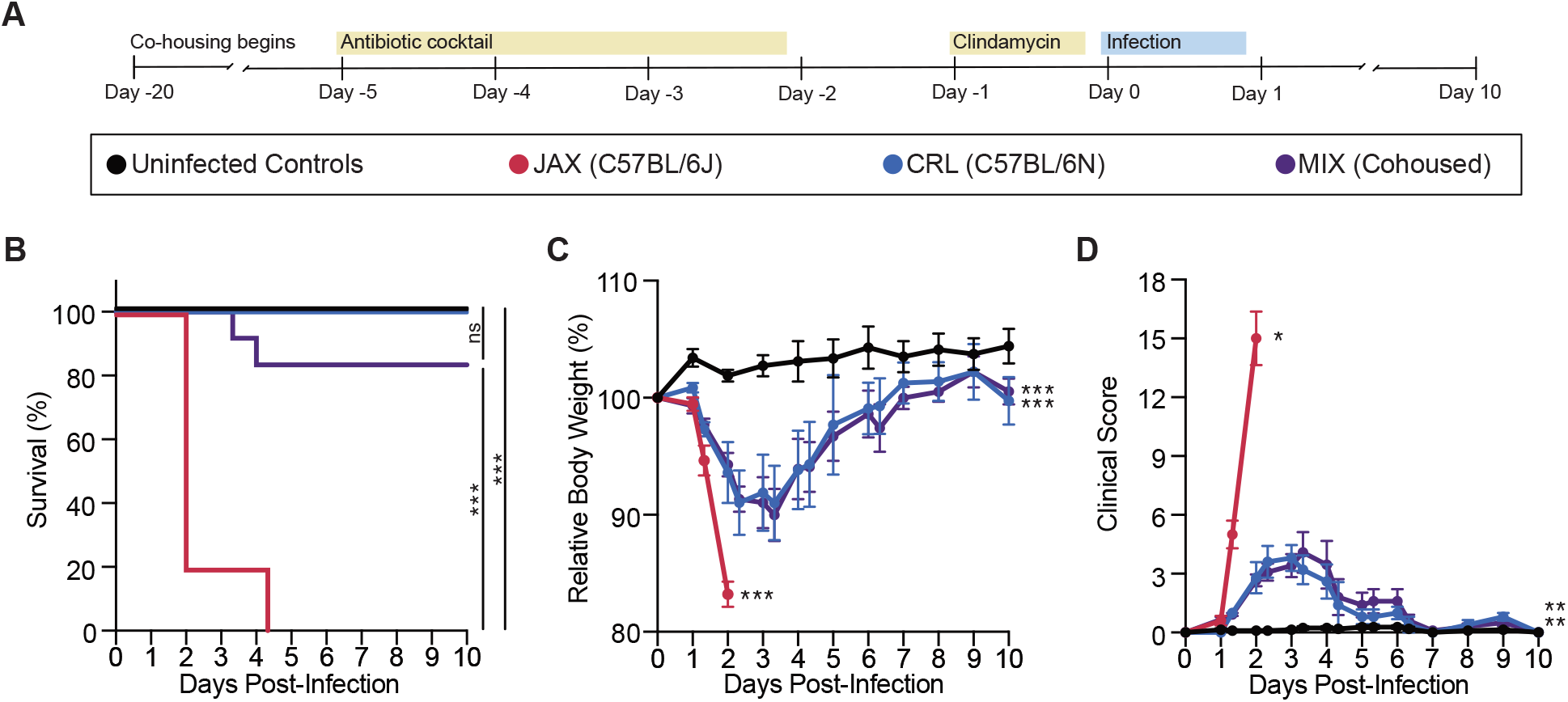
Cohousing modifies disease outcomes following *C. difficile* challenge in JAX and CRL mice. (A) Experimental design: JAX and CRL mice were cohoused (MIX) or housed separately (JAX, CRL) for 14 days, then treated with antibiotics and challenged with either 0 (uninfected controls) or 1x10^6^ CFU spores. (B) Kaplan-Meier survival curve. (C) Relative body weight (% of baseline at Day 0). (D) Disease severity following *C. difficile* challenge. Panels (C) and (D) show mean ± SEM. *ns P* ≥ 0.05, \**P* ≤ 0.05, \*\**P* ≤ 0.01; *\*\*\*P* ≤ 0.001 (comparisons to uninfected control, unless specified). n = 5-12 mice per group (see Fig. S2).

Consistent with our initial findings, separately housed JAX mice exhibited fulminant disease following infection, with all mice succumbing by day 4 post-infection (median survival, 2 days; Figure 2B). In contrast, all separately housed CRL mice survived infection. Notably, cohoused (MIX) mice displayed resistance to infection, with only two deaths observed, one of each vendor origin. Survival in the MIX group did not differ significantly from either the uninfected control or CRL, and was significantly improved relative to JAX mice, indicating that cohousing substantially reduced susceptibility to severe CDI (Figure 2B).

All infected groups exhibited significant weight loss relative to uninfected controls (Figure 2C), as well as clinical signs of disease (Figure 2D; S2A). Fecal *C. difficile* burdens were comparable between CRL and MIX mice, with the same level of colonization and clearance (Figure S2B), indicating that protection in cohoused mice was not attributable to reduced colonization efficiency.

Together, these findings demonstrate that cohousing substantially alters the susceptibility to CDI. While JAX mice housed separately remain highly susceptible to severe disease, cohousing with CRL mice confers significant protection from mortality and severe disease. This transfer of resistance indicates that the different susceptibility observed between vendors is driven, at least in part, by the microbiota.

### JAX and CRL mice harbour distinct microbiota

To determine whether the baseline microbiota of JAX and CRL differed and could explain their different responses to *C. difficile* challenge, fecal samples collected immediately prior to cohousing (Day -20) were analyzed by full-length 16S rRNA gene sequencing (Figure 2A).

To assess alpha diversity, we compared Chao1 richness and Simpson diversity between vendors. Chao1 index was not significantly different between JAX and CRL mice (Figure 3A), whereas Simpson diversity was significantly reduced in JAX mice relative to CRL mice (Figure 3A), indicating a less evenly distributed microbial community in JAX mice at baseline. Beta diversity analysis revealed distinct baseline microbiota between JAX and CRL mice. Principal coordinate analysis based on weighted unique fraction metric (UniFrac) distances showed clear separation between the two vendors (PERMANOVA, *P* = 0.030; Figure 3B), indicating that microbial community structure was associated with vendor origin. Consistent with these findings, relative abundance profiles of genera also reflected clear compositional differences between JAX and CRL mice (Figure 3C).

**FIGURE 3:**
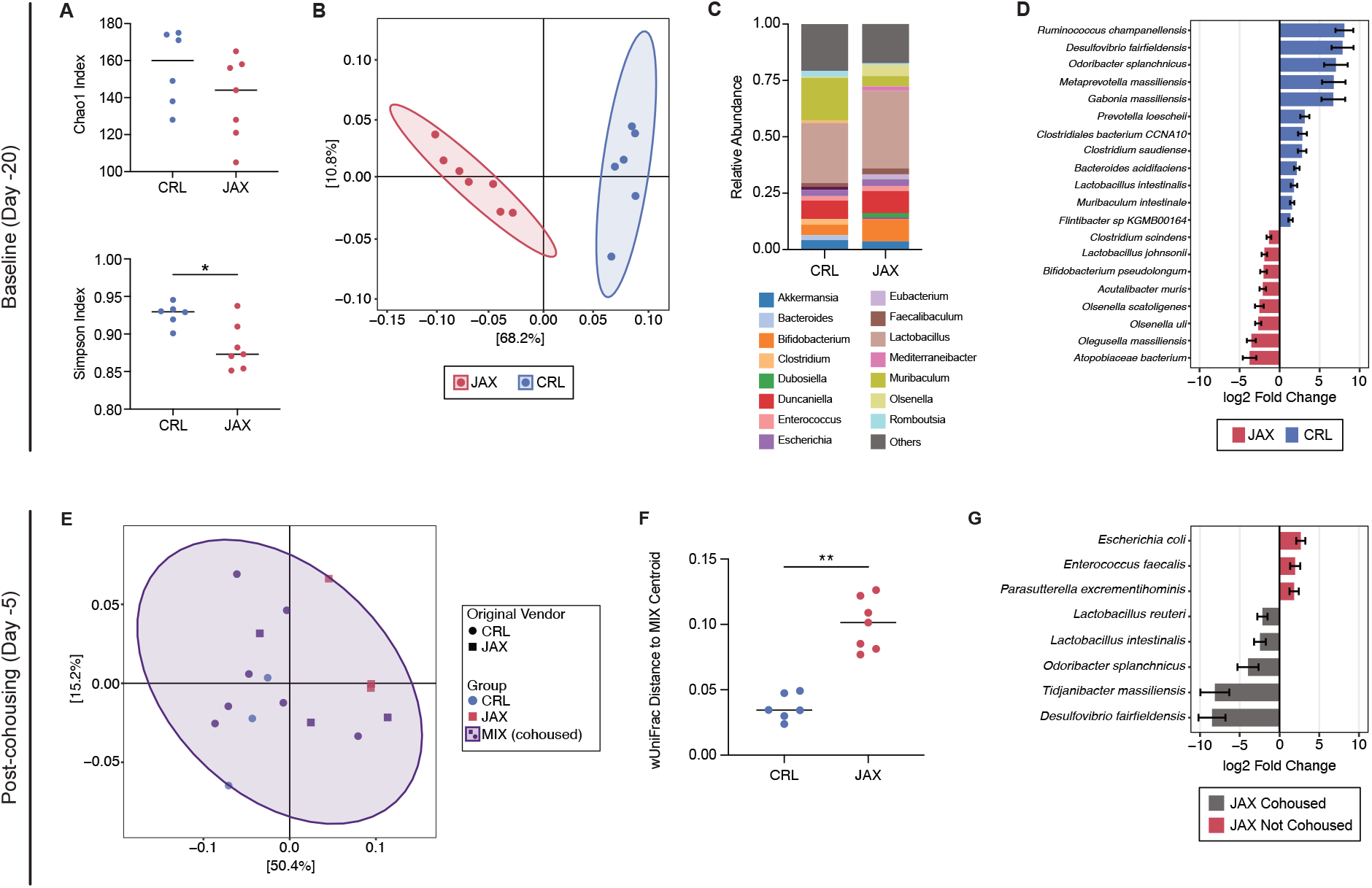
Distinct baseline microbiotas in JAX and CRL mice converge following cohousing. (A-D) Baseline microbiota prior to cohousing (Day -20): (A) Alpha diversity metrics (Chao1 and Simpson indices; Mann-Whitney U Test), (B) principal coordinate analysis (PCoA) of weighted UniFrac distances comparing JAX and CRL mice (PERMANOVA, *P* = 0.030), (C) average relative abundance of the 15 most abundant genera, and (D) differential abundance analysis (DESeq2, FDR ≤ 0.05) showing the 20 most significant species. (E-G) Microbiota after 14 days of cohousing (Day -5): (E) PCoA of weighted UniFrac distances for JAX, CRL, and cohoused (MIX) mice (PERMANOVA), (F) Weighted UniFrac distance from Day -20 CRL and JAX baseline microbiota to the Day -5 MIX centroid (Mann-Whitney U Test) (G) differential abundance analysis (DESeq2, FDR ≤ 0.05) comparing non-cohoused JAX mice to cohoused JAX mice within the MIX group. \**P* ≤ 0.05; \*\**P* ≤ 0.01.

Vendor-dependent differences in baseline microbiota composition were further supported by differential abundance analysis using DESeq2, which identified 59 species that differed significantly between vendors (Figure 3D; Table S1). CRL mice were enriched in several short-chain fatty acid-producing taxa, including *Ruminococcus champanellensis, Odoribacter splanchnicus, Prevotella loeschii, Bacteroides acidifaciens*, and *Muribaculum intestinale*.^33-36^ Additional enrichment was observed for hydrogen sulfide-producing bacterium *Desulfovibrio fairfieldensis*, and lactate-producing *Lactobacillus intestinalis*, among other taxa.^37^ JAX mice were enriched in several Coriobacteriales species including *Atopobiaceae bacterium, Olegusella massiliensis, Olsenella uli*, and *Olsenella scatoligenes*. Additional enrichment was observed of *Bifidobacterium pseudolongum, Lactobacillus johnsonii*, and bile acid metabolizer *Clostridium scindens*.^38^

Together, these results demonstrate that JAX and CRL mice harbour compositionally distinct gut microbiota at baseline.

### Cohousing alters the microbiota of JAX and CRL mice

To assess the impact of cohousing on gut microbiota composition, fecal samples were collected two weeks after cohousing was initiated (Day -5), prior to antibiotic treatment and *C. difficile* challenge (Figure 2A). Sequencing of these samples revealed substantial remodeling of the microbiota following cohousing.

Beta diversity analysis based on weighted UniFrac distances showed that cohoused mice (MIX) exhibited an intermediate community structure relative to JAX and CRL, with MIX mice distributed between the two groups irrespective of their vendor origin (Figure 3E). MIX mice microbiota did not differ significantly from CRL mice (PERMANOVA, *P* = 0.33; Figure 3E), but was distinct from JAX mice (PERMANOVA, *P* = 0.049; Figure 3E), indicating a shift of the MIX community toward the CRL microbiota structure following cohousing. To confirm that this shift reflected cohousing rather than baseline drift, non-cohoused JAX and CRL communities at Day - 5 were compared with their respective Day -20 baselines. No differences were observed for either vendor across timepoints (PERMANOVA: CRL, *P* = 0.23; JAX, *P* = 0.062; Figure S3), indicating that the microbiota remained stable over the 14-day period in the absence of cohousing. To quantify the directionality of this shift, the average weighted UniFrac distance was calculated from Day - 20 JAX and CRL baseline samples to the Day -5 MIX centroid. The average distance from CRL to MIX was significantly lower than from JAX to MIX (Figure 3F), confirming that the MIX microbiota shifted more strongly toward a CRL structure than JAX composition.

Differential abundance analysis by DESeq2 was used to identify vendor-specific compositional shifts induced by cohousing, comparing cohoused and non-cohoused mice within each vendor at Day -5. Remodeling was more extensive in JAX mice than in CRL mice. Cohoused JAX mice showed depletion of *Escherichia coli, Enterococcus faecalis*, and *Parasutterella excrementihominis* relative to non-cohoused JAX mice, alongside enrichment of *Desulfovibrio fairfieldensis, Tidjanibacter massiliensis, Odoribacter splanchnicus, Lactobacillus intestinalis*, and *Lactobacillus reuteri (*Figure 3G; Table S2). In contrast, differentially abundant taxa in cohoused CRL mice were limited to a decrease in *Eubacterium coprostanoligenes* and an increase in *Marvinbryantia formatexigens* relative to non-cohoused CRL mice (Table S3).

Together, these results indicate that cohousing altered the microbiota of both JAX and CRL mice, producing partial convergence with disproportionately greater remodeling occurring in cohoused JAX mice.

### JAX mice exhibit a distinct response to antibiotic treatment

To evaluate how antibiotic treatment reshaped the gut microbiota, fecal samples were collected on the morning of infection, following completion of antibiotic treatment (Day 0; Figure 2A). This sampling point followed a 3-day oral broad-spectrum antibiotic cocktail (kanamycin, gentamicin, colistin, metronidazole, and vancomycin) and a single intraperitoneal dose of clindamycin prior to infection, as described previously.^21^

Alpha diversity analysis revealed significant differences in taxonomic richness between groups. JAX mice displayed significantly lower Chao1 richness compared with CRL and MIX mice, while no difference was observed between CRL and MIX animals (Figure 4A), indicating that antibiotic treatment produced a more pronounced loss of taxonomic richness in JAX mice. In contrast, Simpson diversity was comparable across all three groups (Figure 4A), suggesting that despite differences in richness, the remaining taxa were distributed with similar evenness.

**Figure 4:**
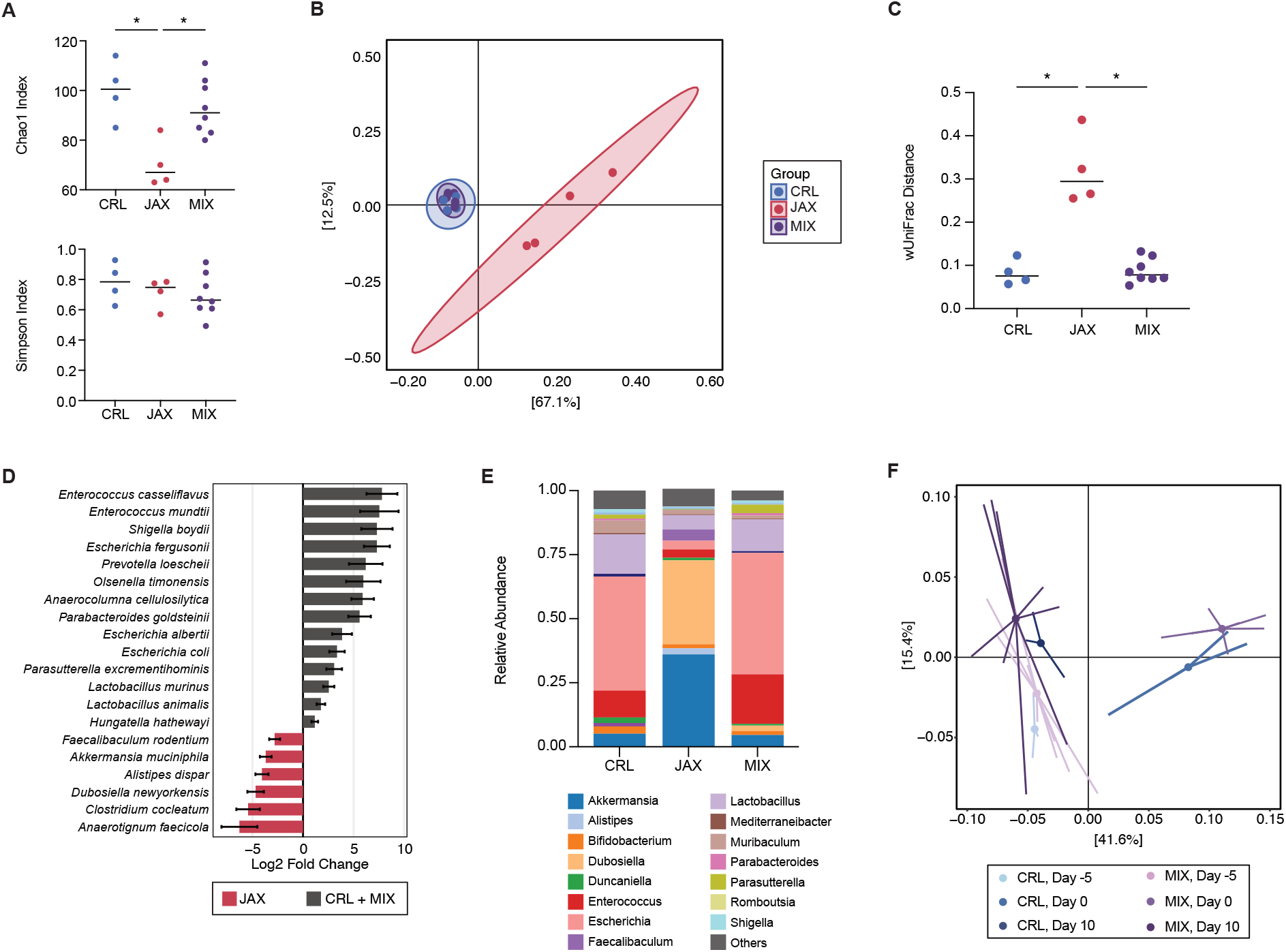
Microbiota at infection and recovery: (A-E) Microbiota at the day of infection (Day 0): (A) Alpha diversity metrics (Chao1 and Simpson indices; Kruskal-Wallis with Dunn’s test) for CRL, JAX, and cohoused (MIX) mice, (B) principal coordinate analysis (PCoA) of weighted UniFrac distances (PERMANOVA: JAX vs. CRL, *P* = 0.025; JAX vs. MIX, *P* = 0.003, CRL vs. MIX, *P* = 0.56) (C) weighted UniFrac distance from CRL, JAX, and MIX samples to the MIX centroid (Kruskal Wallis Test with Dunn’s test), (D) differential abundance analysis (DESeq2, FDR ≤ 0.05) showing the 20 most significant species distinguishing JAX from CRL and MIX mice, (E) average relative abundance of the 15 most abundant genera, and (F) PCoA of weighted UniFrac distances comparing CRL and MIX microbiota at Day -5, Day 0, and Day 10 (PERMANOVA). \**P* ≤ 0.05.

Beta diversity analysis revealed distinct community structures following antibiotic treatment. Weighted UniFrac PCoA showed a clear separation of the JAX microbiota from both CRL and MIX communities (PERMANOVA: JAX vs. CRL, *P* = 0.025; JAX vs. MIX, *P* = 0.003; Figure 4B), with no significant difference observed between CRL and MIX mice (*P* = 0.56; Figure 4B). To further quantify this separation, the dispersion of each group relative to the MIX centroid was assessed. JAX samples were significantly more dispersed from the MIX centroid than either CRL or MIX samples, while CRL and MIX samples did not differ from one another (Figure 4C). Together, these confirm that antibiotic treatment drove a pronounced and distinct restructuring of the JAX microbiota relative to cohoused and CRL mice.

To identify taxa distinguishing these two communities, DESeq2 was used to compare JAX mice with CRL and MIX mice (pooled CRL and MIX). CRL and MIX mice were enriched in several members of Pseudomonadota (*Shigella boydii, Escherichia albertii, Escherichia coli), Enterococcus* spp. (*E. casseliflavus, E. mundtii)*, and *Lactobacillus* spp. (*L. murinus, L. animalis)*, among other taxa (Figure 4D; Table S4). In contrast, JAX mice were enriched in *Akkermansia muciniphila* and *Dubosiella newyorkensis*, along with *Alistipes dispar, Faecalibaculum rodentium, Clostridium cocleatum*, and *Anaerotignum* (Figure 4D; Table S4). Consistent with these findings, relative abundance profiles of dominant genera confirmed that *Akkermansia* and *Dubosiella* together accounted for a disproportionately large fraction of the JAX microbiota at Day 0 (Figure 4E).

Together, these findings indicate that antibiotic treatment produced vendor-dependent shifts in microbiota composition. CRL and MIX mice retained taxonomically enriched and compositionally similar communities, whereas JAX mice developed a compositionally distinct and depleted microbiota. These differences immediately preceding infection are consistent with the divergent susceptibility to *C. difficile* challenge subsequently observed between groups.

### Cohousing-associated microbiota convergence persists following *C. difficile* infection

To determine whether microbiota convergence persisted after infection, fecal samples were collected from surviving mice at day 10 post-infection. Survivors at this timepoint consisted exclusively of CRL and MIX mice. Weighted UniFrac PCoA was performed across three timepoints spanning post-cohousing (Day -5), post-antibiotic treatment (Day 0), and post-infection recovery (Day 10). At Day 10, CRL and MIX survivors exhibited indistinguishable community structures (PERMANOVA, *P* = 0.47; Figure 4F), demonstrating that microbiota convergence between groups was maintained through antibiotic treatment and recovery from infection. CRL mice showed no significant difference between their Day 10 and Day -5 community structures (PERMANOVA, *P* = 0.10; Figure 4F), indicating recovery toward a pre-antibiotic community composition. Together, these results demonstrate that the MIX microbiota converged toward, and remained aligned with, the CRL community structure throughout antibiotic treatment and *C. difficile* challenge.

Consistent with these community-level findings, DESeq2 differential abundance analysis identified only a limited number of taxa distinguishing CRL and MIX survivors at Day 10 (Table S5). MIX survivors were enriched for several *Alistipes* species (*A. onderdonkii, A. putredinis, A. finegoldii, A. senegalensis*, and *A. shahii*), as well as *Anaerotignum faecicicola* and *Hungateiclostridium thermocellum*, while CRL survivors were enriched for *Enterocloster bolteae, Lachnoclostridium* sp. *YL32*, and *Clostridiales bacterium CCNA10* (Table S5). Despite these differences, the overall similarity in community structure between CRL and MIX survivors at this timepoint indicates that cohousing restructuring of the microbiota was largely preserved throughout the course of infection and recovery.

## DISCUSSION

In this study, we demonstrate that the outcome following *C. difficile* challenge is vendor dependent. While JAX mice were susceptible to fulminant disease, CRL mice were resistant. This difference was driven by the microbiota rather than host genetics, as the resistant CRL phenotype was transferable to susceptible JAX mice via cohousing, with cohoused JAX mice converging toward the CRL microbiota structure. These findings extend prior work establishing the microbiota as a key determinant of CDI severity,^39, 40^ and parallel a *Salmonella typhimurium* model in which cohousing C57BL/6J with C57BL/6N mice reduced mortality via transfer of protective *Enterobacteriaceae*.^30^ Together, these results establish the microbiota as the primary determinant of CDI outcome in this model, and motivates closer examination of the specific taxa and functional pathways which distinguishes the resistant CRL community from its susceptible JAX counterpart.

The use of cohousing in this experiment exploits the coprophagic behaviour of mice to enable spontaneous and bidirectional microbial exchange.^41^ The resulting exchange was asymmetric, as MIX mice clustered more closely with CRL than JAX mice before and after antibiotic treatment, indicating that the CRL microbiota dominated, rather than blended, with the JAX community. Although JAX and CRL represent distinct substrains with known differences in physiology,^28^ the magnitude of phenotype convergence here suggests that these host-level differences are subordinate to the microbiota-mediated effects on infection outcome.

Vendor-dependent differences in susceptibility to CDI have been reported previously, attributed to colonization of mice with the toxin-positive but low-virulence *C. difficile* strain LEM1.^42^ In our study, *C. difficile* was not detectable in uninfected control mice from either vendor following antibiotic treatment, ruling out an endogenous bloom of *C. difficile* as the source of protection in CRL mice. This indicates that the vendor-associated disease severity is driven by differences in the resident gut microbiota rather than pre-existing colonization with a non-virulent *C. difficile* strain.

This study relied on longitudinal 16S rRNA gene sequencing, which provided taxonomic resolution of the microbiota in response to cohousing, antibiotic treatment, and infection. However, these findings cannot provide functional resolution of the microbiota, therefore the specific taxa highlighted below should be interpreted as candidate contributors rather than confirmed drivers of protection or susceptibility.

At baseline, CRL and JAX mice harboured distinct communities, with CRL being more taxonomically even, in addition to being enriched in several SCFA-producing taxa compared to JAX (Figure 3B). SCFA-producing taxa are critical to protection against CDI, by reinforcing the integrity of the colon epithelial barrier^10, 43^, immunomodulation and ensuring the correct activation of the inflammasome in the colonic epithelium^44^, and by directly antagonizing *C. difficile* outgrowth.^45, 46^

Following cohousing, *O. splanchnicus* became enriched in cohoused JAX mice compared to non-cohoused JAX mice, consistent with transfer from the CRL microbiota (Figure 3G). Notably, *O. splanchnicus* has been associated with decreased risk of *C. difficile* acquisition in a clinical cohort.^47^ Mechanistically, *O. splanchnicus* ferments butyrate, which restricts *C. difficile* outgrowth and toxin production, and has been shown to increase the pool of secondary bile acids in a colitis mouse model.^48^ More broadly, *O. splanchnicus* is considered a core member of a healthy gut microbiota, with reported immunomodulatory effects including the expansion of regulatory T cells and IL-10 production, and protection against *Salmonella typhimurium* and *Listeria monocytogenes* mouse infection models.^49, 50^ Together, *O. splanchnicus* is a plausible contributor to the milder disease phenotype observed in CRL and cohoused JAX mice. Simultaneously, cohousing resulted in a depletion of *E. faecalis* in cohoused JAX mice (Figure 3G). Although this preceded infection, *E. faecalis* has been shown to exacerbate *C. difficile* fitness and pathogenesis by cross-feeding fermentable amino acids,^51^ suggesting its downregulation may have contributed to a less permissive environment for *C. difficile* in cohoused JAX mice.

Following antibiotic treatment, JAX mice underwent a more severe contraction of taxonomic richness than CRL or MIX mice (Figure 4A), resulting in a community dominated by *A. muciniphila* and *D. newyorkensis* (Figure 4D; 4E). *A. muciniphila* is a mucin-degrading specialist, that uses glycoside hydrolases to liberate glycan oligosaccharides from host mucus, which serve as carbon and nitrogen sources for itself and neighbouring bacteria.^52^ While mucin degradation by *A. muciniphila* is generally important for maintaining mucosal homeostasis, unchecked expansion can erode the mucus layer, exposing the underlying epithelium and increasing susceptibility to bacterial translocation, toxin-mediated damage, and inflammation.^53^ Consistent with this, antibiotic-induced expansion of *A. muciniphila* has been shown to thin the mucus layer, damage goblet cells, and induce inflammation^54^, and similarly increases susceptibility to *Citrobacter rodentium*.^55^ Notably, *A. muciniphila* can directly crossfeed *C. difficile* by releasing mucin-derived monosaccharides from MUC2.^52^ Together, these suggest that antibiotic-induced expansion of *A. muciniphila* in JAX mice may provide a nutrient source supporting *C. difficile* colonization and compromise mucosal integrity, predisposing the epithelium to toxin-mediated damage and inflammation during infection.

In contrast, CRL and MIX mice retained a more taxonomically enriched community at the time of infection (Figure 4A). Interestingly, despite resilience against disease, these communities were dominated by Pseudomonadota. Expansion of Pseudomonadota, particularly *Escherichia-Shigella*, is a typical signature of clindamycin-induced dysbiosis in the mouse microbiota,^56-58^ and expansion of these taxa are associated with colonization and symptomatic *C. difficile* in humans.^59, 60^

The changes in community composition over time suggest that resistance and susceptibility to CDI are unlikely to be attributable to any single bacteria but rather reflect the overall structure of the microbiota and its fluctuation throughout antibiotic treatment and infection. Targeted metabolomic profiling of SCFAs and secondary bile acids, both of which directly influence *C. difficile* outgrowth and virulence^2^, would help clarify whether these taxonomic differences translate to functionally relevant differences in metabolite availability. Additionally, small sample sizes may have limited the detection of subtle compositional differences, and the use of only two vendors leaves it unclear whether the taxa identified here generalize beyond this specific pair, or importantly into the context of a human microbiota. Future studies incorporating metabolomics, larger cohorts, and additional microbiota sources would help to further resolve the functional basis of the microbiota-mediated resistance to CDI.

These findings demonstrate that vendor-associated differences in gut microbiota composition determine susceptibility to *C. difficile* infection, and that susceptibility can be transferred between hosts through cohousing. The composition of the microbiota is an important and overlooked source of variability in preclinical CDI models, underscoring the need to account for vendor origin and baseline microbiota composition when designing and interpreting mouse studies of CDI. Identifying the specific taxa and metabolic pathways that confer this protection may inform future microbiota-targeted strategies, such as defined bacterial consortia, for CDI prevention and treatment.

## METHODS

### Strain and Culturing Conditions

*Clostridioides difficile* strain VPI 10463 (ATCC 42355) was used for *C. difficile* challenge experiments. All culturing was performed at 37 ºC in a vinyl anaerobic chamber (Coy Laboratories, Grass Lake, MI, USA), maintained with a gas mixture of 3% H_2_, 10% CO_2_, 87% N_2_.

### Spore Preparation

Spores were prepared as described previously, with minor modifications.^61^ Briefly, a single colony of *C. difficile* was inoculated into brain heart infusion (BHI) broth (BD Difco, Franklin, NJ, USA) and incubated for 48h. Cultures (200 µL) were spread onto 70:30 sporulation agar plates and incubated anaerobically at 37 ºC for 3 weeks to allow sporulation. Plates were flooded with 3 mL sterile water, and spores was harvested by scraping with a sterile loop. The suspension was centrifuged at 21 000 x g for 10 min, the supernatant discarded, and the pellet was resuspended in sterile water. Suspensions were stored at 4 ºC for 3 days and subsequently suspended in lysozyme solution (2 mg/mL lysozyme in 10 mM Tris-HCl, pH 8.0, containing 1% (v/v) Triton X-100) for 2 h at 37ºC. Samples were sonicated and purified by centrifugation through a 50% (w/v) sucrose gradient. The spore pellet was collected, washed repeatedly with sterile water, and stored at 4 ºC until use. Spore preparations were enumerated using a hemocytometer under phase-contrast microscopy.

### Mice

Female C57BL/6 mice (5-7 weeks old) mice were obtained from The Jackson Laboratory (Bar Harbor, ME, USA) and from Charles River Laboratories (St Constant, QC, Canada). Mice were housed under specific pathogen-free conditions in the Comparative Medicine and Animal Resources Centre at McGill University. All animal procedures were approved by the Animal Care Committee of McGill University under the animal use protocol MCGL-10104 and conducted in accordance with guidelines from the Canadian Council on Animal Care.

#### (1) *Clostridioides difficile* challenge

Beginning 5 days prior to infection, mice received an antibiotic cocktail for 3 days consisting of kanamycin (80 mg/kg; Thermo Fisher Scientific, Waltham, MA, USA), gentamicin (7 mg/kg; Sigma-Aldrich, St Louis, MO, USA), colistin (8.4 mg/kg; Sigma-Aldrich), metronidazole (43 mg/kg; Sigma-Aldrich), and vancomycin (8.4 mg/kg; VWR, Radnor, PA, USA). Antibiotics were administered three times daily in 20 µL sterile drinking water by micropipette-guided oral administration, as described previously, to ensure consistent dosing between animals.^62^ One day prior to infection, mice received clindamycin (10 mg/kg; MilliporeSigma, Burlington, MA, USA) via intraperitoneal injection. Mice were challenged by oral gavage with 10^5^, 10^6^, or 10^7^ spores in 100 µL sterile PBS. Mice were monitored three times daily and scored for weight loss, activity, posture, coat condition, diarrhea, and ocular/nasal symptoms, as previously described.^63^ Mice reaching ≥ 20% weight loss or score ≥ 12/18 were euthanized according to humane endpoint criteria. Animals that died prior to euthanasia were assigned a maximal clinical score of 18.

#### (2) Cohousing Experiment

For the cohousing experiment, C57BL/6J (JAX) and C57BL/6N (CRL) mice were either housed separately with animals from the same vendor or cohoused in groups of four mice per cage (two JAX and two CRL mice). Cohousing began 2 weeks prior to antibiotic treatment and *C. difficile* challenge to allow microbiota transfer via coprophagy.^41^ Mice were administered antibiotics and infected with 10^6^ spores as described above.

### Quantification of *C. difficile* Colonization

Fecal samples were collected and homogenized in anaerobically reduced PBS to generate a 10% w/v suspension. Insoluble material was removed by centrifugation at 50 x g for 30 s. The fecal slurry was serially diluted and plated onto Brucella agar (MilliporeSigma) supplemented with *Clostridium difficile* selective supplement (Oxoid, Basingstoke, UK) and 0.1% taurocholic acid, as previously described.^64^ Plates were incubated anaerobically at 37ºC for 48 h prior to colony enumeration.

### DNA Extraction

Fecal samples were collected throughout the experiment and stored at -80 ºC until processing. Microbial DNA was extracted using the ZymoBIOMICS DNA Miniprep Kit (Zymo Research, Irvine, CA, USA) following the manufacturer’s instructions.

### 16S rRNA gene Sequencing

16S rRNA gene sequencing was performed as described previously.^65^ Briefly, 16S amplicons were generated by combining 2X Platinum SuperFi II Green PCR Master Mix (Invitrogen, Carlsbad, CA, USA), custom M13-27F and M13-1492R barcoded primers (0.2 µM; Table S6), and 2 µL of template (∼ 25 ng) in a reaction volume of 25 µL. Thermocycling was performed with initial denaturation at 98 ºC for 30 s, followed by 15 cycles of 98 ºC for 10s, 60 ºC for 10 s, and 72 ºC for 45 s, followed by a final extension at 72 ºC for 5 min. Amplicon concentration was quantified with the Qubit dsDNA HS Assay Kit (Invitrogen) and pooled at equal concentrations. Pooled amplicons were cleaned using AMPure XP beads (Beckman Coulter, Brea, CA, USA), washed with 80% ethanol, and eluted in 28 µL nuclease-free water. 2 µL of the pool was tagged with DBCO by combining with 2X Platinum SuperFi II Green PCR Master Mix and DBCO-M13 primer (0.5 µM). Thermocycling conditions were identical to the initial PCR but with only 10 cycles. The tagged pool was cleaned using AMPure XP beads as described above and eluted in 25 µL elution buffer (Oxford Nanopore Technologies, Oxford, UK). The sequencing library was prepared by combining 11 µL of eluted DNA with 0.3 µL Rapid Adapter and 0.7 µL Adapter Buffer, and incubating at 37 ºC for 10 min. 12 µL of the prepared library was combined with 37.5 µL sequencing buffer and 25.5 µL library beads, then loaded onto a primed Flongle flow cell (Oxford Nanopore, Version R10.4.1) in a MinION device, following the SQK-RAD114 Rapid Sequencing V14 protocol. Sequencing was performed overnight. Basecalling was performed using Dorado (v0.8.3) using the SUP super high-accuracy model. Reads were filtered for lengths of 1500 ± 200 bp, then assigned to taxa using Emu (v3.4.4, https://github.com/treangenlab/emu)^66^. Counts were imported into R (v2025.05.1+513) as a phyloseq object (https://github.com/joey711/phyloseq)^67^. Alpha and beta diversity analyses were performed using vegan (v2.7.3 https://github.com/vegandevs/vegan)^68^, and ape (v5.8.1 https://github.com/emmanuelparadis/ape)^69^. Differential abundance analyses and relative abundance were conducted using MicrobiomeAnalyst (v3.0, https://www.microbiomeanalyst.ca/)^70^.

### Statistical Analysis & Graphing

Statistical methods were not used to determine sample sizes, and the investigators were not blinded to experimental groups. Survival was analyzed using the Mantel-Cox method with Holm-Bonferroni correction for multiple comparisons. Weight and clinical scores were excluded from analysis when fewer than three animals remained per group. Weight loss was compared using two-way ANOVA, followed by Tukey’s multiple comparison test. Average clinical scores were compared using Kruskal-Wallis followed by Benjamini-Hochberg procedure. Alpha diversity metrics were compared by the Mann-Whitney U Test or by Kruskal-Wallis followed by Benjamini-Hochberg procedure. Beta diversity was assessed by PERMANOVA using the *pairwiseAdonis* package (v0.4.1, https://github.com/pmartinezarbizu/pairwiseAdonis)^71^. Details for each statistical test are provided in the figure legends. Data were plotted in GraphPad Prism (v11.0.0) or with *ggplot2* (v4.0.2, https://github.com/tidyverse/ggplot2)^72^, and figures were assembled in Affinity Designer 2 (v2.6.5).

## Data Availability

16S rRNA gene sequencing reads were deposited in NCBI BioProject ID PRJNA1515306.

## ACKNOWLEDGEMENTS

We thank the staff of the Comparative Medicine and Animal Resource Centre for their contributions to the animal experiments. L.K. is the recipient of a Canadian Institutes of Health Research (CIHR) Canada Graduate Research Scholarship and a Fonds de Recherche du Québec Doctoral award. This work was supported by CIHR project grants (PJT-173262 and PJT-203764) to B.C.

## SUPPLEMENTARY FIGURES

**Supplementary Figure 1:**
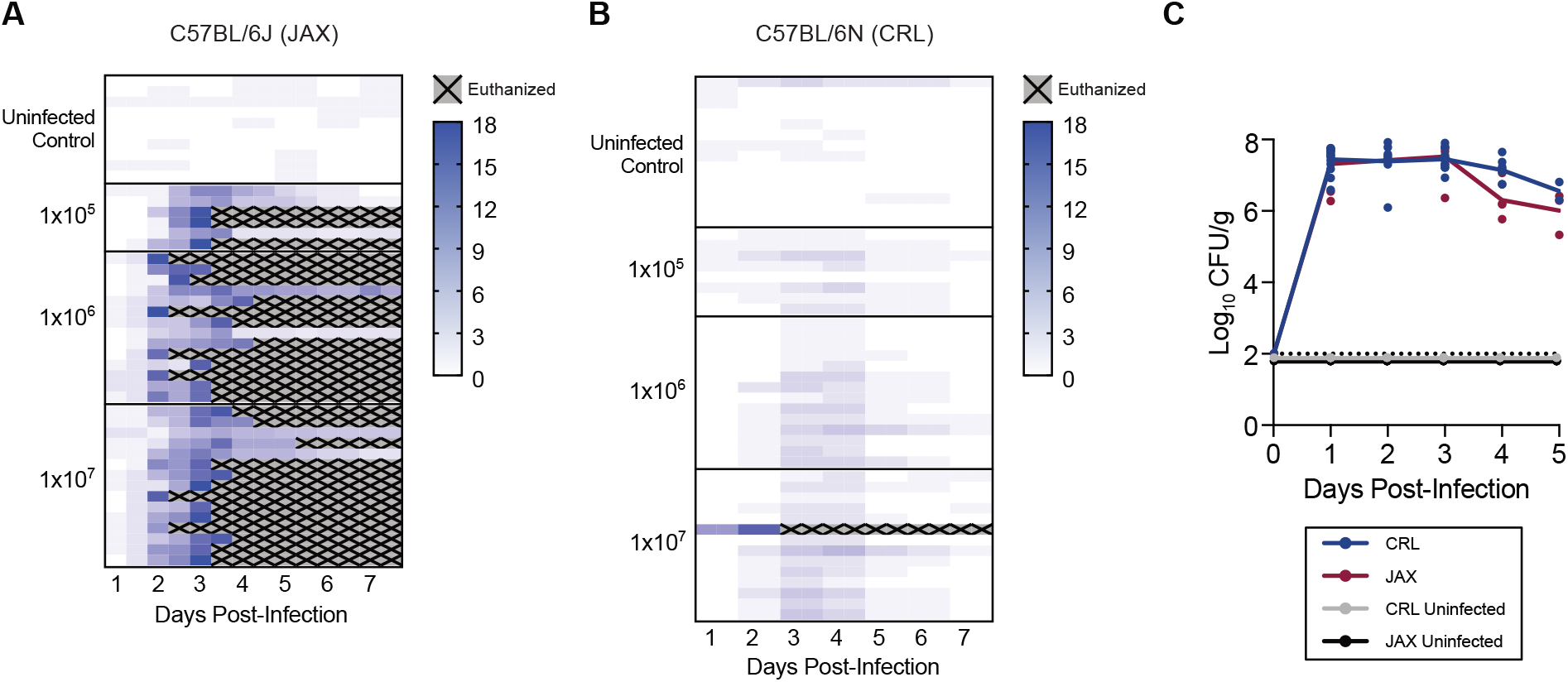
Individual disease severity scores and *C. difficile* colonization following infection. (A) JAX mice and (B) CRL mice individual disease severity scores grouped by *C. difficile* inoculum. (C) *C. difficile* colonization levels (Log_10_CFU/g feces) in fecal samples from JAX and CRL mice post-infection with inoculum 1x10^7^ CFU. No significant differences in colonization between vendors (JAX vs. CRL) (2-way ANOVA, vendor effect *P* = 0.057). Dashed line indicates the limit of detection (100 CFU).

**Supplementary Figure 2:**
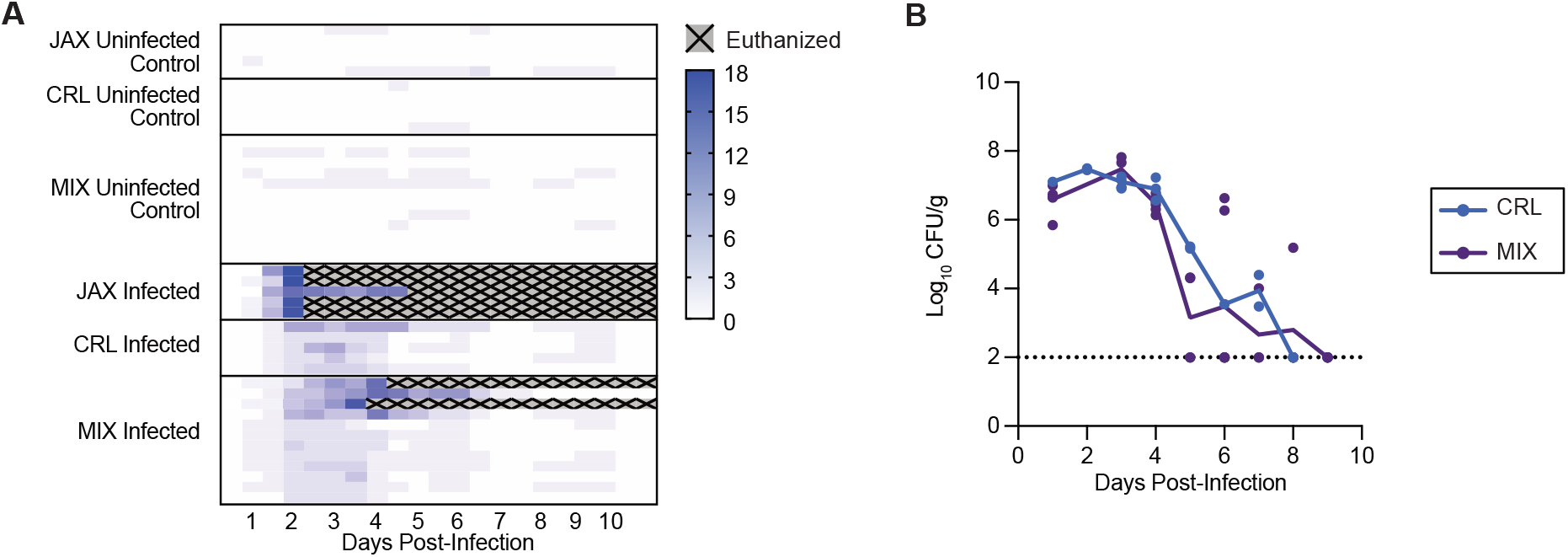
Individual disease severity scores and *C. difficile* colonization following infection in cohousing experiment. (A) Individual clinical scores of cohoused (MIX) and separately housed (JAX, CRL) mice following *C. difficile* challenge. (B) *C. difficile* colonization levels (CFU/g feces) in fecal samples collected from cohoused (MIX) and separately housed (CRL) mice post-infection with 1x10^6^ CFU spores. No significant difference in colonization between CRL and MIX groups (two-way ANOVA, *P* = 0.53). Fecal samples could not be collected from JAX mice post-infection. Dashed line indicates the limit of detection (LOD = 100 CFU).

**Supplementary Figure 3:**
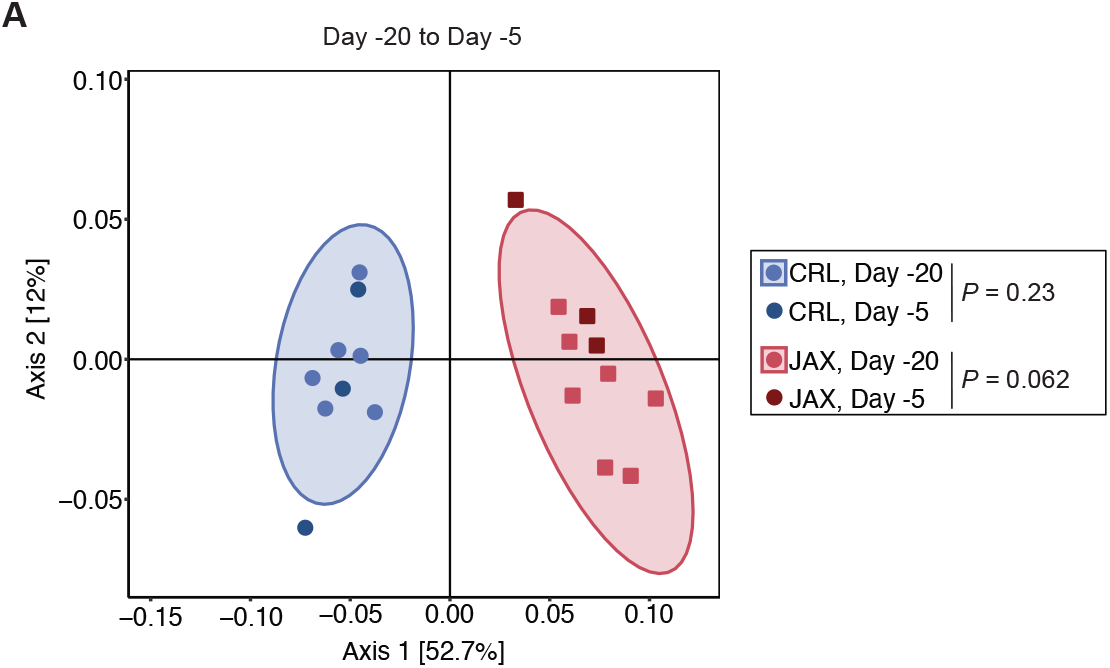
Microbiota remains stable over time in the absence of cohousing. (A) PCoA of weighted UniFrac distances for separately housed CRL and JAX mice at Day -20 (baseline) and Day -5 (concurrent timepoint during the cohousing period). PERMANOVA was used to test within-vendor community stability across timepoints. (CRL *P* = 0.23, JAX *P* = 0.062).

